# The Readiness Potential reflects expectation, not uncertainty, in the timing of action

**DOI:** 10.1101/2020.04.16.045344

**Authors:** Eoin Travers, Maja Friedemann, Patrick Haggard

**Affiliations:** Institute of Cognitive Neuroscience University College London; Department of Experimental Psychology University of Oxford

## Abstract

Actions are guided by a combination of external cues, internal intentions and stored knowledge. Self-initiated *voluntary actions*, produced without any immediate external cue, may be preceded by a slow EEG Readiness Potential (RP) that progressively increases prior to action. The cognitive significance of this neural event is controversial. Some accounts link the RP to the fact that timing of voluntary actions is generated endogenously, without external constraints, and perhaps even randomly. Other accounts take the RP as reflecting the unique role of planning, therefore of temporal expectation, in voluntary actions. In many previous experiments, actions are both unconstrained by external cues, but also potentially involve preplanning and anticipation. To separate these factors, we developed a reinforcement learning paradigm where participants learned, through trial and error, the optimal time to act. If the RP reflects freedom from external constraint, its amplitude should be greater early in learning, when participants do not yet know the best time to act. Conversely, if the RP reflects planning, it should be greater later on, when participants have learned, and know in advance, the time of action. We found that RP amplitudes grew with learning, suggesting that this neural activity reflects planning and anticipation for the forthcoming action, rather than freedom from external constraint.

## Introduction

Human actions are guided by a combination of external cues, internal intentions and stored knowledge. Self-initiated *voluntary actions* – those produced without any immediate external prompt – are preceded by a slow EEG Readiness Potential (RP) that ramps up over the second or so prior to action (Deecke et al., 1969). The RP is primarily generated by the Supplementary and pre-Supplementary Motor Areas (SMA and pre-SMA; Shibasaki & Hallett, 2006), which in turn receive strong drive from the subcortical circuitry of the basal ganglia. Interestingly, the readiness potential is reduced or absent prior to externally-triggered actions that are temporally matched to each participant’s voluntary actions (Jenkins et al., 2000). This result suggests that it is a specific neural correlate of voluntary action.

Why does the RP occur, and what does it represent? Accounts of the RP tend to focus on one of two contrasting facets of self-initiated actions. Some accounts emphasise the random, unconstrained, and unpredictable nature of self-initiated actions (Eccles, 1985; Jo, Hinterberger, Wittmann, Borghardt, & Schmidt, 2013; Nachev, Rees, Parton, Kennard, & Husain, 2005; Schurger, Sitt, & Dehaene, 2012). Others emphasise the unique role of planning and temporal expectations in internally-generated actions (Brunia, Boxtel, & Böcker, 2011; Verleger, Haake, Baur, & Śmigasiewicz, 2016).

### Randomness and Uncertainty in Action

Schurger and colleagues (Erra, Arbotto, & Schurger, 2019; Schurger, Sitt, & Dehaene, 2012; see also Murakami, Shteingart, Loewenstein, & Mainen, 2017) recently proposed that the timing of self-initiated actions may effectively be random. In this model, actions are triggered when randomly fluctuating neural activity reaches a threshold. When this activity is time-locked to action and averaged across trials – the usual way of analysing EEG from RP experiments – this model reproduces the classic time course of the RP. We will refer to this account as the *stochastic model*. The RP usually begins a second or more prior to action (Shibasaki & Hallett, 2006). When participants are asked when they decided to move, they typically indicate a time just moments prior to action (Libet, 1985), leading to the interesting conclusion that voluntary actions must have unconscious causes (Libet et al, 1983). The stochastic model explains this apparent contradiction. It holds that there is no specific neural event corresponds to the start of the RP, and the apparent “onset” of of the RP is instead an artefact of averaging across trials. Any decision to move would in fact occur after a stochastic neural signal reaches some predefined threshold for action.

Could self-initiated actions really be random? Random actions can sometimes be useful (Glimcher, 2005; Maye et al., 2007). In animals, the ability to produce actions that cannot be predicted by prey or by predators can have an adaptive value (Brembs, 2011; Maye et al., 2007; Maynard Smith, 1982). Randomness also plays an important role in optimal models of decision-making in uncertain environments. In a familiar environment, the optimal policy is to consistently choose whatever action has the best pay-off. In an uncertain environment, one must strike a balance between exploiting immediately available options and exploring potentially better alternatives (March, 1991). A common way to achieve this is through stochastic action selection: usually choosing what one believes to be the best option, but sometimes randomly exploring other options instead (Gershman, 2018). The Thompson sampling algorithm (Thompson, 1933), where the degree of randomness is proportional to the agent’s uncertainty about which action is best, is close to the optimal policy in many environments, and approximates human behaviour (Gershman, 2018).

Nachev and colleagues (Nachev et al., 2005, 2008) proposed a slightly different view. They argued that SMA and pre-SMA activity, and hence presumably the RP, reflects conflict or uncertainty due to the lack of external constraint on self-initiated actions. In simple self-initiated actions, conflict occurs because there are many possible times at which one could act, but no reason to favour one over the others – the decision to act is *underdetermined*. Nachev et al. (2008) therefore suggest that medial frontal activity, as measured by RP, may reflect uncertainty about *when* to act, just as the anterior cingulate, located just ventral to SMA, reflects uncertainty about *which* action to perform (Botvinick et al., 2004). This account, like the stochastic account, predicts a greater RP when the reasons for action are least clear or most uncertain.

Zapparoli et al. (2018; see also Seghezzi et al., 2019) report MRI results consistent with this idea. They asked participants to perform actions in response to cues, or to freely decide either what action to perform, when to act, and whether or not to go ahead with the action (Brass & Haggard, 2008). They found stronger SMA activation for free actions than cued actions, and found that SMA activity was stronger for free decisions about when to act than for free decisions about what action to perform, or whether to act.

### Planning, Temporal Expectation, and Action

Other explanations link the RP not to randomness, but to determination and imposition of structure and pattern on human behaviour. These views emphasise the unique role of planning and temporal expectation in self-initiated actions. The sensory consequences of a voluntary action are thought to be automatically and unconsciously predicted by a *forward model* (Blakemore et al., 2002). Thus, self-initiated actions are events that the actor expects to occur (Friston et al., 2010), and which represent a reduction in entropy, or the imposition of an internal model on the external world (Parr & Friston, 2019). This has two implications. First, an agent can begin to prepare an action long before they intend to perform it. Second, they can have expectations about when an action and its consequences will occur. The RP could reflect either of these processes.

Voluntary movements must be prepared before they are executed (Lara et al., 2018; Wise, 1985). Direct motor and premotor cortical recording from monkeys and rodents (e.g. Elsayed, Lara, Kaufman, Churchland, & Cunningham, 2016; Lara et al., 2018; Murakami et al., 2017) show that motor preparation involves passing through a sequence of neural states. Similar preparatory states may occur prior to self-initiated actions and prior to speeded, externally-triggered actions, but preparation of self-paced actions often takes more time (Lara et al., 2018). The RP reflects the firing rates of populations of motor and supplementary motor neurons (Shibasaki & Hallett, 2006). Therefore, it provides a one-dimensional readout of the multidimensional pre-movement neural activity captured by direct cortical recordings. This implies that the RP may not be a distinctive feature of self-initiated actions. Instead, the RP could reflect a general process of motor preparation, which can either be performed rapidly in response to an imperative stimulus, or, if necessary, initiated endogenously, and extended over a long period of time.

There is also a strong link between motor preparation and temporal expectation. People often prepare in advance to ensure they can act at the right time. The medial frontal cortex, including SMA, is also involved in temporal anticipation of future events (Tecce, 1972) and in processing the passage of time (Kolling & O’Reilly, 2018). The RP is strikingly similar to the Contingent Negative Variation (CNV; Brunia et al., 2011; Tecce, 1972), a slow negative component that ramps up prior to the time at which participants expect to receive a behaviourally relevant stimulus. CNV is typically recorded in reaction time experiments where a warning cue occurs at a fixed interval prior to a “Go” signal. A number of authors have argued that the RP and CNV both reflect slow motor preparation (Brunia et al., 2011; Grünewald et al., 1979; Rohrbaugh & Gaillard, 1983): the RP represents preparation of a self-timed action, while CNV represents preparation of an action timed to a predictable external event. This conclusion is supported by source localisation of CNV to supplementary motor areas (Hultin et al., 1996). Interestingly, a smaller CNV occurs even when participants are not required to respond to the stimulus, suggesting that it also reflects non-motor temporal processes (Rohrbaugh & Gaillard, 1983). The RP may also occur in the absence of immediate action, since it is apparently also found prior to “covert decisions” that do not immediately lead to actions (Alexander et al., 2016; Gluth et al., 2013). These findings indicate that the RP and CNV may reflect both motor preparation and temporal expectation.

The contribution of temporal expectation to the RP is supported by classical findings. Libet and colleagues (Libet, 1985; Libet et al., 1983) distinguished between Type I and Type II RPs (see also Frith & Haggard, 2018). When participants preplanned in advance the time at which they would move, RPs began early and reached a high amplitude (Type II RPs). When participants instead acted “freely and capriciously” (Libet et al., 1983) without extensive preplanning, RPs began just before action, and reached lower amplitudes (Type I RPs). However, the distinction between Type I and Type II RPs was made on the basis of participants’ subjective reports about their general strategy for initiating actions. These reports were apparently made at the end of a block of several trials, so cannot be readily linked to individual action events. Other studies (e.g.Schurger, 2018) have noted that the RP tends to be greater on trials where participants spontaneously waited longer than usual before acting.

### Randomness or Planning?

In most studies of the RP, both planning and expectation, and randomness and uncertainty, might all play a role in action. Participants may produce actions that are unconstrained by external stimuli, and so appear random and unpredictable to an observer. Nonetheless, these actions might well be preplanned, and so the movement and its consequences may be expected by the participant themselves. In other words, random behaviour might in fact be carefully planned.

As a result, these descriptive studies cannot determine whether the RP reflects a process of planning and predetermination, or randomness and indetermination. To separate these two possible influences, we need an experimental design which explicitly manipulates the degree of determination/indeterminacy of action, and investigates how this affects the RP. Here we do this by providing a context where participants must learn, through trial and error, the optimal time to act. This design encourages gradual acquisition of preparatory planning for action, as participants learn when they should act. The task thus encourages a progressive learning-related shift from “capricious”, random behaviour to regular, preplanned behaviour (Libet, 1985). Our design would initially encourage random exploration of the environment, for example, by acting at various different times, and monitoring the outcome. These early actions are unpredictable and unconstrained by the environment. While such exploratory actions might in principle be preplanned, preplanning them offers no obvious advantage over simply relying on a random generator. Later, after becoming more certain about when they should act, participants should consistently wait a fixed time before doing so. These later, post-learning actions would be strongly constrained by the environment, or at least by the historical environment of previous actions and outcomes. There is an obvious advantage to preplanning them, and an obvious disadvantage to allowing their timing to vary randomly. Therefore, learning the optimal time to act offers an experimentally-tractable way to investigate whether the RP reflects the process of determining an action, or rather represents the indeterminacy and randomness of action generation (figure 1). If the RP reflects randomness and absence of constraint, RPs should decrease over time as participants progress from random exploration towards planning based on an internal model of the optimal time interval. If the RP instead reflects planning, RPs should increase over with learning (if we assume that initial exploratory actions are not preplanned but late actions are preplanned) or should at least stay constant (if we assume that even initial exploratory actions in fact derive from strategic plans to sample across the distribution of action times).

**Figure 1.**
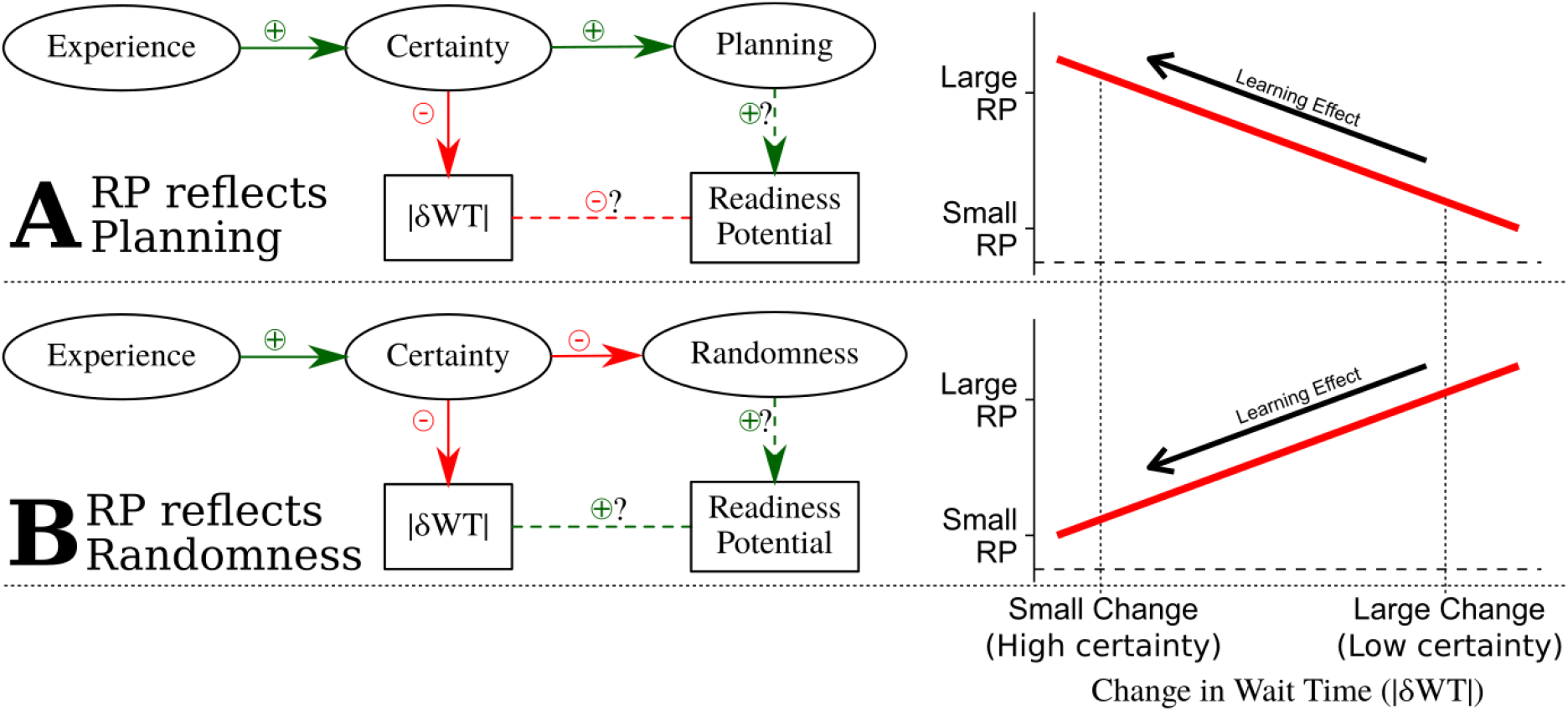
Predictions. Ovals show latent variables, rectangles observed variables, and arrows causal effects. Question marks indicate hypothesised relationships. As participants gain experience over the course of each block, their certainty in the correct time to act increases, trial-to-trial changes in their waiting times (|*δWT*|) decrease, planning, certainty and temporal expectation increase, and the role of randomness or stochasticity and uncertainty in action timing decreases. **A.** If planning drives the RP, it will have greatest amplitude on trials with small changes in waiting times (negative relationship between |*δWT*| and RP magnitude). **B.** If randomness or uncertainty drive the RP, it will be greatest on trials with large changes in waiting times (positive relationship between |*δWT*| and RP magnitude). Since our learning task required reduction of temporal uncertainty, we show the effects of learning not as a function of time or trial number, but as our behavioural proxy for (un)certainty, namely the trial-to-trial update in action time (|*δWT*|) that indicates progressive acquisition of the to-be-learned optimal time of action.

## Method

### Participants

We aimed to test 20 participants, based on the sample size used for previous RP studies (e.g. Khalighinejad et al., 2018). Twenty-one healthy participants (11 female, mean age ± SD: 24.24 ± 4.37 years) took part in the experiment. We excluded two participants who moved excessively during EEG recording. Thus, 19 participants were included in the analyses (10 female, mean age ± SD: 23.53 ± 3.58 years). Participants received £7.50 per hour reimbursement upon completing the experiment, plus a bonus for performance on the task. The experiment was approved by the UCL ICN ethics committee and each participant’s written informed consent was collected before starting the experiment.

### Procedure

We developed a temporal reinforcement learning paradigm that allows participants to learn, through experience, the best time to act. Our cover story treated participants as bakers, placing a soufflé in an oven at the beginning of each trial. Their task was to wait until the soufflé was ready, and then press a key to withdraw it from the oven to gain a small cash bonus. If participants withdrew a soufflé before it was ready, that trial was aborted and the next began after a short interval. Thus, participants learned the optimal baking/waiting time through this action feedback. Crucially, the average baking/waiting time required varied between blocks, and had to be learned from feedback.

Participants completed 15 blocks in total, taking 3 minutes each, and were instructed to score as many points as possible in that time. Soufflés were ready after an average of 3, 5, 7, 9, or 11 seconds depending on the block, with a SD of 1 second across trials in each block. There was no explicit penalty for leaving the soufflé too long, but doing so reduced the number of trials a participant could complete in the time available.

Before starting the experiment, participants were shown two sample trials by the experimenter. Following this, participants completed one three-minute round in order to get familiar with the task and to check their understanding. The baking time of soufflés in the trial round was normally distributed with a standard deviation of 1 and a mean of 7, giving the participants a reasonable prior expectation for the real task.

If participants knew the distribution of baking times for each block, the optimal strategy would be to wait for time τ when they are sufficiently certain that the soufflé was ready before withdrawing it. Since they had to learn this information, the optimal strategy is to initially explore the effects of opening the oven (that is, acting) after different times to estimate τ, and then to use that estimate to score as many points as possible. This can be achieved through Bayesian learning strategies such as Thompson sampling. An agent using such a strategy begins each block with low posterior precision in their estimate of τ. They will thus be highly stochastic in their actions, but learn quickly from new evidence. As they learn, their estimate becomes more precise, their actions less stochastic, and their response to new evidence reduced.

### EEG Acquisition

The experiment was conducted in an electrically shielded room. We used a BioSemi ActiveTwo system (BioSemi, 2011) to record 32-channel EEG. Electrodes were placed in locations FP1, FP2, F7, F3, Fz, F4, F8, FC5, FC1, FCz, FC2, FC6, T7, C3, C1, Cz, C2, C4, T8, CP5, CP1, CPz, CP2, CP6, P7, P3, Pz, P4, P8, O1, Oz, and O2. This montage includes a higher than usual density of electrodes clustered around Cz to capture motor activity. Electro-oculogram (EOG) was recorded with electrodes above and below the right eye and on the outer canthi of both eyes to control for eye movement artefacts. EEG was sampled at 500 Hz.

### EEG Preprocessing

EEG data preprocessing was performed with Python using the MNE software package (Gramfort et al., 2013). Data were bandpass filtered between 0.1 and 250 Hz, and downsampled to 250 Hz for analysis. As electrodes were more densely clustered around Cz, a pure average reference would disproportionately subtract activity from this area. Ideally, an average reference signal should give equal weights to signals from all areas of the scalp. Therefore, we used the average of all electrodes excluding FCz, C1, C2, CPz, T7, and T8 as a more representative reference signal. Data with large motor artefacts were removed by visual inspection. Independent component analysis (ICA) was then used to identify eye movement and blink artefacts from the EEG data. Eye-movement related ICA components were identified by visual inspection and by correlating their activity with that of the EOG channels.

RP epochs were extracted from 3 seconds before to 0.5 seconds after action. Epoch recordings were baseline-corrected to the average of the window −3 to −2.9 seconds prior to action. Epochs with amplitude values exceeding 60 μV from baseline were excluded from the analysis. To reduce noise for single-trial analyses, we averaged the signal from the five electrodes around location FCz: Fz, FC1, FCz, FC2, and Cz. To quantify the RP amplitude on a single trial, we compared the average voltage on these electrodes in the final 50 ms prior to action, to a baseline defined as the average signal between 3.1 and 3 s prior to action (Figure 3B).

## Results

### Behavioural results

Participants’ behaviour is consistent with a simple iterative reinforcement learning strategy (Figure 2B-C). If their action was too early (i.e., feedback showed soufflé not yet ready) on trial *t-1*, participants waited 1.9 s (SD = 0.7 s) longer on trial t, t(19) = t(19) = 12.63, p < .001. If their action was later than required (i.e., feedback confirmed that soufflé was already baked on trial *t-1*), participants acted 0.4 s (SD = 0.4 s) sooner on trial t, t(19) = 4.28, p < .001.

**Figure 2.**
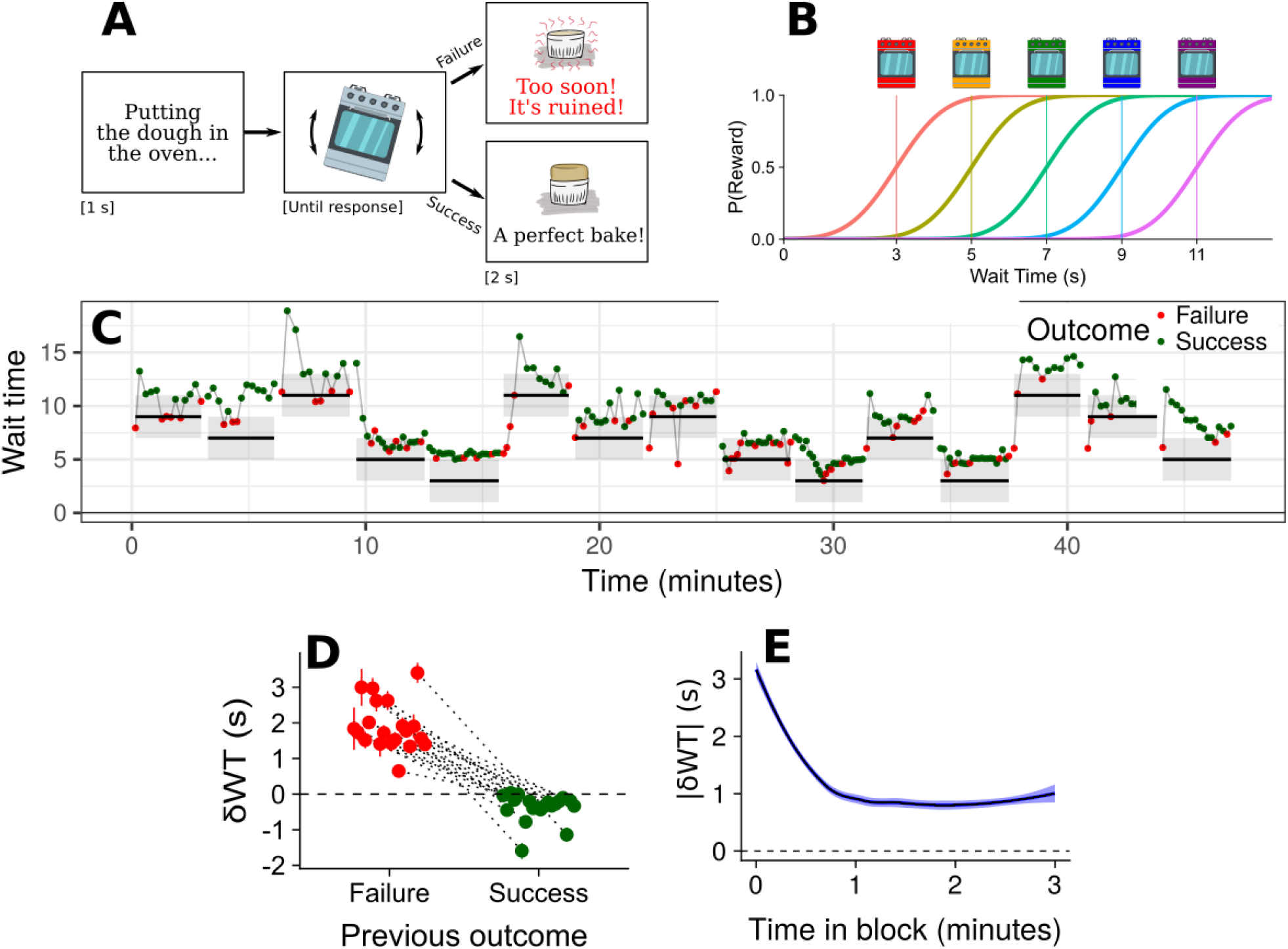
Temporal reinforcement-learning task and behavioural results. **A.** Outline of behavioural task. A soufflé was placed in the oven at the start of each trial, and participants had to wait an unknown time before pressing a button to withdraw it. Points were awarded only on trials where action was not premature. **B.** The time participants needed to wait before acting differed across conditions (differed coloured ovens), and had to be learned through experience. **C.** Waiting times across the experiment for a single participant. Horizontal bars show mean ±2SD times at which soufflés were ready in each block. Green (red) dots show trials where participants did (not) wait long enough. **D.** Participants waited considerably longer after trials where they acted too soon (failures) and acted slightly sooner after a previous trial where they waited long enough to retrieve the soufflé (successes). **E.** Trial-to-trial changes in waiting times decreased over the course of each block. Thus, people learned a better time for action.

The amount participants adjusted their wait times in response to feedback decreased progressively in each block (Figure 2C), indicating strong learning initially, and weaker learning later on. By reducing their learning rates in this way, participants can home in on the optimal waiting time in a way that approximates Bayesian learning. In other words, participants should, and do, substantially update how long they wait before acting in response to feedback early in learning, when they are uncertain of the correct time. Conversely, later in learning they should update their wait times only slightly, because they have acquired greater certainty about the optimal time to wait. We used absolute, unsigned changes in participants’ wait time from one trial to the next, |δWT|, as a measure of their uncertainty about the optimal time to act, and so of how exploratory or as a measure of their uncertainty about the optimal time to act, and so of how exploratory or stochastic their actions were. This lets us characterize individual responses as being more exploratory (strongly different from time of previous action, therefore high |δWT| or more exploitative (repeating time of action from previous trial, therefore low |δWT|).

We use |δWT| rather than trial number, or progress through a block, as our proxy for certainty in later analyses (see also figure 1). We do so for several reasons. First, the number of trials a participant completes per block depends on how long they wait on each trial. Trial counts therefore vary considerably between participants and between blocks, and trial number cannot readily be compared across blocks. Second, the time and number of trials needed to converge on a consistent waiting time varies between participants and blocks, so that a participant might be very certain about when to act after one minute, or after 10 trials, on one block, but be unsure at the same point in time, or after the same number of trials, in another block. Since |δWT| is a consequence of participants’ level of certainty, which is the focus of the learning task, it avoids these ambiguities. Further, on some trials participants updated their waiting times in the wrong direction, acting later after success or sooner after premature responses. We assume that the absolute magnitude of the update indicates that they were uncertain. In contrast, the *sign* of the update may simply indicate whether they responded appropriately to their own uncertainty. We therefore used the absolute magnitude of the update in waiting time in either direction as our measure of certainty. A small update in either direction is taken to reflect high certainty about when to act, and a large update is taken to reflect high uncertainty.

### EEG Analysis

Analyses were conducted using the lme4 package for R (Bates et al., 2015). To test the effect of uncertainty while controlling for possible confounds, we fit a linear mixed model with single-trial RP voltage as the dependent variable, |δWT| as our predictor of interest, and signed δWT, WT, and block number as covariates. All predictors were centred on their mean value within participants. Random intercept terms were included for each participant. More complex random effects structures yielded singular variance-covariance matrices, and so could not be interpreted. For visualisation purposes (Figure 3C), we coded trials as being either above the median value of |δWT| for that block (*exploratory responses*) or below it (*exploitative responses*). Since |δWT| is compared to the median value within each particular block, half of the trials in each block are coded as exploratory, half as exploitative. This eliminates any confounds due to systematic differences between blocks.

**Figure 3.**
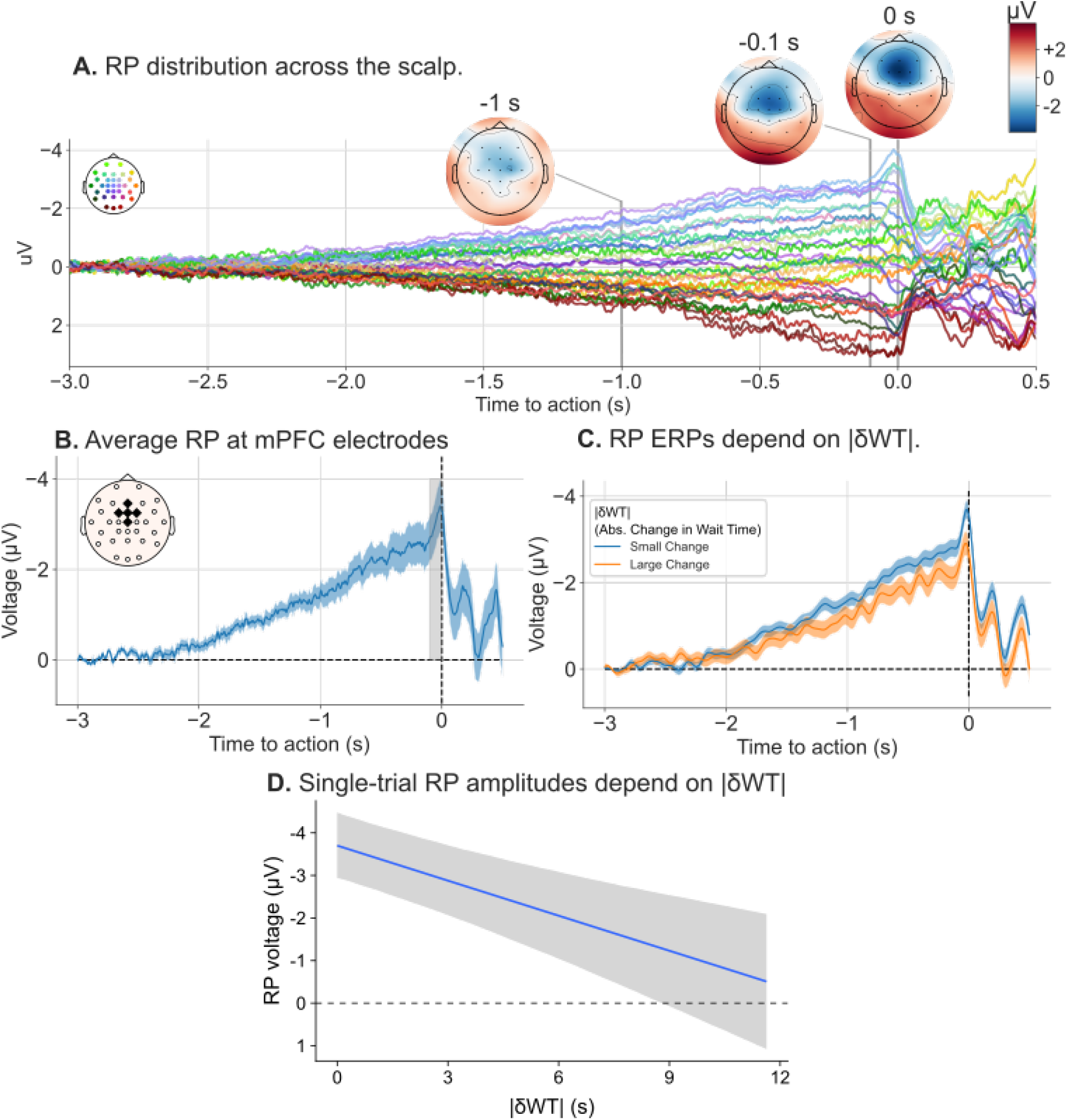
EEG Results. **A.** A clear Readiness Potential (RP) occurred prior to action. **B.** RP Event-Related-Potential (ERP) averaged across centro-frontal electrodes (Fz, FC1, FCz, FC2, and Cz). The average voltage in the last 50 ms prior to action (grey) was used as an estimate of single-trial RP voltages. **C.** RP ERPs for actions where absolute changes in wait times (|δWT|) were larger (orange) or smaller (blue) than the median for that block. **D.** Estimate (±SE) of single-trial RP voltages as a function of |δWT| after adjusting for other terms in the model.

### EEG Results

A clear RP occurred prior to participants’ responses in this task (Figure 3A-B). Average RP ERPs for explore and exploit responses are plotted in Figure 3C. The RP appears to have a greater amplitude explore actions (high |δWT|) than exploit actions (low |δWT|) throughout most of its time course. This was confirmed by the linear mixed model analysis on RP amplitudes in the final phase before action. Thus, there was a significant effect of |δWT| on RP amplitudes, b = 0.28 CI = [0.04, 0.53], t(3308.1) = −2.240, p = .030, meaning that greater trial-to-trial updates in waiting times were associated with lower (less negative) RPs. This is consistent with the averaged ERP results. There was no effect of signed δWT| (increasing or decreasing wait times), B = 0.03 CI = [−0.16, 0.22], t(3307.9) = 0.280, p = 0.780, of actual wait time (longer or shorter wait times), B = −0.03 CI = [−0.15, 0.09], t(3307.4) = −0.540, p = .590, or of block number, B = −0.01 CI = [−0.09, 0.06], t(3309.8) = −0.380, p = .700.

## Discussion

Does the Readiness Potential reflect randomness and uncertainty, or planning and expectation? We found that RP amplitude increases as participants learn through experience how long to wait before acting, so that their certainty about action time increases, and their actions become less random, more preplanned and more predictable. This is consistent with the proposal that the RP reflects anticipatory planning and preparation to make an action at a specific time in the future. It appears inconsistent, however, with the proposal that the RP reflects uncertainty (Nachev et al., 2008), or arises as the result of a purely stochastic triggering process (Schurger et al., 2012).

### Accumulator Models of Action

Schurger et al. (2012) proposed that spontaneous self-initiated actions could be triggered by a neural circuit that accumulates random noise until it reaches a threshold. They showed that the RP could reflect random fluctuations in accumulated noise, time-locked to the time they crossed threshold. How might such a model apply to our paradigm?

One possibility is that the Schurger et al. (2012) model applies only to pure self-initiated actions where timing is unspecified, and not to actions during model-based cognitive tasks such as ours, where there is an optimal time to act. However, there was a clear RP prior to action in our task (Figure 3A). If this model does not apply here, we must conclude that it is not an explanation of the RP in general, but rather an explanation of the RP under specific condition that participants are asked to act, but given no guidance at all about when to act.

Second, actions might be generated by several different pathways, only some of which give rise to an RP. Thus, the lateral premotor pathway for externally-triggered actions has been distinguished from the medial frontal pathway, based on pre-SMA and SMA, for internally-triggered actions (Passingham, 1993). Similarly, there might be one mechanism for generating spontaneous, arbitrary actions, and one for deliberate, preplanned movements. Our results might arise because the latter generates stronger RPs than the former. This view recalls Libet et al.’s (1983) distinction between Type I and Type II RPs. An analogous distinction was proposed by Maoz et al. (2019), who reported a smaller RP prior to responses to value-based decisions (choosing to donate to one of two possible charities) than prior to arbitrary actions (pressing one of two buttons at random). Those authors suggested that arbitrary actions are triggered by the accumulation of noise in SMA (Schurger et al., 2012), leading to an RP, while value-based actions are triggered by another mechanism, possibly in ventromedial prefrontal cortex, that does not lead to an RP (Wallis, 2007).

In most contexts, including our task, actions are neither purely random nor purely value-based, but are hybrid actions driven by some mixture of these processes (Luce, 1959). Any particular action might be triggered by just one of these mechanisms (e.g., Obhi & Haggard, 2004), or by a combination of the two (see Hughes et al., 2011). In any event, the random mechanism may be assumed to play a greater role earlier compared to later in learning, while the converse holds for the value-based mechanism. If the random accumulation mechanism is the generator of the RP, we would expect to find an RP of greater amplitude early in learning. Since we found the opposite effect, our results are inconsistent with the interpretation of the RP as reflecting a random process for triggering actions, at least in the context of the present task.

A third possibility is that the same neural accumulation mechanism is responsible for the timing of both random, arbitrary actions and planned, value-based actions, but that the input to the accumulator differs. Arbitrary actions could be triggered by random noise in the accumulator, while value-based actions are triggered by a specific external input. We have simulated this possibility.

Briefly, the RP can indeed be reproduced by a model that accumulates random noise in the absence of specific inputs. However, we find that the same model predicts greater RP amplitudes when driven by clear evidence that one should *act now*. We illustrate this idea in Figure 4, and describe the simulations used to reach it below. The full simulation parameters and the python code used to conduct the simulations can be found in Supplementary Materials.

**Figure 4.**
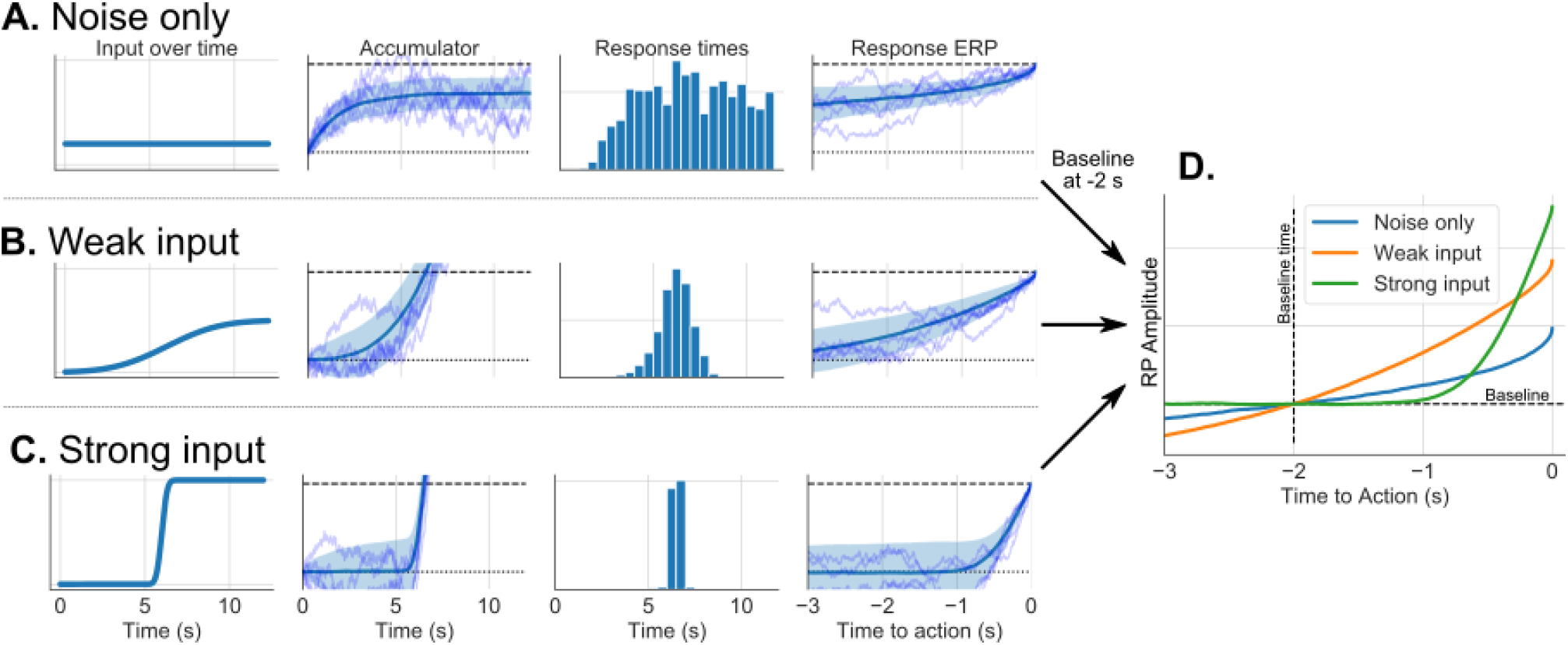
Simulation results. Simulations from a model where **A.**) the observer is highly uncertain of the correct time to act (constant weak input;, **B.**) somewhat uncertain (weak gradually increasing input;, or **C.**)is highly certain (strong, temporally precise input. **D.** Simulated RP amplitudes, baseline-corrected 2 s before action. Although the threshold for action is the same, differences in activity at the (arbitrarily-chosen) baseline time mean RP amplitudes appears greatest when action is triggered by a strong input signal, and weakest when triggered by stochastic noise.

In our simulations we assume that the input *I_t_* to the accumulator over time depends on the agents’ posterior belief that an action at that time would be rewarded. Since the distribution of reward times is Gaussian, the posterior estimate of P(Reward|Act Now), marginalising over possible readiness times *μ*, is a cumulative Gaussian function centred around 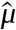, an estimate of the average time taken for a soufflé to be ready, with a slope β that increases as the estimate of 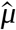 becomes more precise: 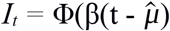, where Φ is the cumulative Gaussian function. Prior to any experience, uncertainty about 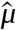 is extremely high, so β is close to 0, the input to the accumulator is weak and constant over time, and the timing of action is largely determined by random noise in the accumulator (Figure 4A). This is equivalent to the model proposed by Schurger et al. (2012), As the estimate of 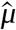 becomes more precise, the input to the accumulator becomes more temporally specific, and actions are triggered directly by this input (Figure 4B-C). As a result, simulated wait times cluster more closely around a time shortly after 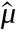.

What happens to the RP? In these models, actions are triggered once the accumulator reaches a fixed threshold. This means that the level of the accumulator at the time of action is the same in all cases. However, the state of the accumulator during the baseline window a few seconds prior to action can change across simulations, and these changes can account for differences in the apparent amplitude of the RP. Therefore, we will focus on the shape of the simulated RP, and the time at which it appears to begin rising to threshold. When actions are driven by noise (Figure 4A), the simulated RP obtained by time-locking accumulator traces to the simulated response times reproduces the classic shape of the RP (Schurger et al, 2012). Since the shape of the RP here is an artefact of biased sampling of random fluctuations, there is no specific moment at which the RP begins. Instead, the slope is steepest close to the time of action, and appears progressively shallower further back in time. This also means that the accumulator is already moderately activated 2 s prior to action. As a result, the amplitude of the RP – the change in activity from −2 s to the time of action – is small here.

As participants become more certain of the optimal time of action (Figure 4B), there is a clear input to the accumulator prior to actions, and the simulated RPs have a clearer onset time. Later still, when the correct time to act is precisely known (Figure 4C), the input to the model ramps up sharply at this time, and the accumulator quickly rises to threshold. From these simulations, we can see that the apparent amplitude of the RP depends not on the state of the accumulator at the time of action, but on how much the accumulator changes between the baseline and the time of action. As a result, the apparent RP amplitude is largest when actions are triggered by a strong input signal, and smallest when triggered by stochastic fluctuations in the accumulator. Thus, if the RP wholly or partly reflects stochastic fluctuations in an evidence accumulation process, then increasing the contribution of the stochastic process does not necessarily produce a larger RP. Rather, greater stochasticity leads to a slower, more gradual rise of the RP.

These results highlight a fundamental limitation of EEG and related approaches. These recordings allow us to infer changes in neural activity over time, but not absolute firing rates. Thus, we can only estimate the amount of activity at the time of action relative to a pre-action baseline, and not the actual level of activity at the time of action. We can try to reduce the risk of distorting apparent RP amplitudes by using a baseline window long before action and a longer ERP. Unfortunately, EEG data contains high-amplitude, low-frequency noise components, and the high-pass filter used to attenuate this noise would also distort longer ERPs. These issues are avoided by direct electrophysiological recordings, although these are only rarely possible in humans. Notably, direct recordings in macaques (Lara et al, 2018) show that the same patterns of firing rates occur prior to self-initiated and externally-triggered actions, consistent with the idea that there is a constant accumulator threshold for action.

### Readiness Potentials and Anticipation

An important limitation of these accumulator models is that they do not capture the kind of temporal expectation and anticipation that some claim the RP reflects (e.g. Brunia et al., 2011). In an accumulator model the state of the accumulator will either ramp up quickly if there is a strong external signal, or ramp up slowly if the external signal is weak or absent. In contrast, temporal expectation can be captured in predictive processing models of cognition and motor control (Blakemore et al., 2002; Wolpert et al., 2003). These models are commonly used to explain the *consequences* of self-initiated actions (Farrer & Frith, 2002; Haggard, 2017), but do not generally focus on the *precursors* of actions, nor on temporal dynamics in the seconds *prior* to action. Thus, they cannot readily provide a mechanistic explanation of preparation and RP. A challenge for future work will be to reconcile evidence accumulation and predictive processing accounts. To do this, it will be necessary to develop models that capture the temporal dynamics of evidence accumulation both during action preparation and during action outcome representation.

Some authors argue that RP and CNV both reflect a single underlying process of temporal anticipation and motor preparation (Brunia et al., 2011; Rohrbaugh & Gaillard, 1983). Our results are consistent with this theory. However, another interpretation is possible. It might be that the RP reflects spontaneity and uncertainty in self-generated actions, while CNV reflects temporal anticipation and prediction. Furthermore, it might be that as our participants become more certain about the best time to act, an RP component becomes weaker but a CNV component becomes stronger. Measured EEG would reflect the sum of these components, which cannot be separately identified. It is difficult to rule out this possibility. Although CNV and RP are traditionally studied using different experimental paradigms, they are not otherwise readily distinguishable (Brunia et al., 2011). Consistent with this, we compared scalp topographies for high certainty and low certainty actions in an exploratory analysis, and found no other notable differences beyond the greater peak amplitude around FCz for high certainty actions.

#### Why is motor preparation so slow?

Motor actions, even relatively complex reaching and grasping actions can be initiated within less than 200 milliseconds and completed accurately, and motor preparation takes only some of that time (Lara et al., 2018). The simple keypress required in RP studies should in principle require even less time to prepare. Neural computations are usually metabolically efficient (Hasenstaub et al., 2010). Why does the nervous system expand so much energy maintaining preparatory activity for so long? We have three hypotheses. First, although movement precision is not important in standard RP tasks, it is crucial in other contexts. By preparing movements well in advance whenever possible, the motor system may give itself time to correct for any inaccuracies before movement begins (Churchland & Shenoy, 2007). Second, slow motor preparation may leave time for upcoming actions to be modified or vetoed. This can happen if external factors mean that an action is no longer appropriate (Schultze-Kraft et al., 2016), or if internal feedback from predictive processing indicates that the action will have undesirable consequences. Third, while rapid changes in neural activity can occur, it may be that these abrupt changes are themselves metabolically costly. Thus, slow motor preparation might strike a balance between avoiding unnecessary prolonged periods of elevated firing rates and avoiding overly abrupt changes in firing rates.

### Learning When to Act

We presented participants with a novel temporal decision-making task. While there exists a large body of work on how humans and animals decide *what* action to produce (Bogacz, 2007; Edwards, 1954; O’Connell, Shadlen, Wong-Lin, & Kelly, 2018), and how they learn about the value of alternative actions (Lee, Seo, & Jung, 2012), less research focusses on deciding *when* to produce an action. Even then, work on decisions about when to act has almost exclusively focused on when agents cease sampling sensory evidence, and commit to a decision (Cisek, Puskas, & El-Murr, 2009; Drugowitsch, Moreno-Bote, Churchland, Shadlen, & Pouget, 2012; Ratcliff, 1978). In contrast, our task is a *pure* timing task, in that one must decide when to perform an action, and there is no interaction between the *when* decision and the amount of evidence available to support the decision. In our task, the decision when to act is based on a model learned from previous experience, rather than on current sensory input. In fact, pure timing decisions of this kind are common in natural behaviour, and can be of vital importance. For instance, animals must decide how long to rest between other activities such as foraging or hunting, and prey must decide how long to avoid an area after seeing a predator there. When two animals meet in confrontation, a stand-off often follows. Each animal must then compute when to attack, or run away etc. Human agents must decide not only when to remove food from the oven, but also when to change job, apply for promotion, start a family etc.

Traditionally, computational accounts of temporal decision-making have fallen into one of two categories. Some treat timing decisions as a series of discrete decisions about what to do, for instance deciding once per trial whether to continue pumping up a balloon, or not (Lejuez et al., 2002), Others treat temporal decision-making as simply a prior decision about how long to wait before acting (e.g. Misirlisoy & Haggard, 2013). Here, we treat it as a continuous-time decision about *when* to act. Doing so opens up a whole swathe of new questions. For instance, studies of continuous-space reinforcement learning – decisions about where to act – have shown that humans use a sophisticated and near-optimal approach to generalise across space (Wu, Schulz, Speekenbrink, Nelson, & Meder, 2018). Moving through time is not like moving through space, since we move through time in only one direction, and at a constant rate. The computations underlying temporal decisions remain unclear, although models of neural timing are widespread (Paton & Buonomano, 2018).

Finally, an important topic for future research will be to outline just how endogenous and exogenous causes of action interact in naturalistic decision-making. One promising approach is to consider how participants adjust the amount of external evidence they require before they commit to a decision (Bogacz et al., 2010; Cisek et al., 2009; Ratcliff, 1978). This is commonly captured by decision-models that include an endogenous *urgency signal* (Cisek et al., 2009) or by time-varying decision criteria (Bogacz et al., 2010). Urgency can increase as a function of elapsed time on a single trial. Self-initiated actions such as those required in our task might be caused by an urgency signal in the absence of any external evidence. Mechanisms such as these are likely to play an important role in any theory of endogenous and exogenous action.

## Conclusion

Participants learned through trial and error when to make a simple action. As participants grew more certain about when to act, and became less variable and stochastic in the timing of their actions, the readiness potential prior to their actions became larger in amplitude. This is consistent with the proposal that the RP reflects motor planning or temporal expectation. It is harder to reconcile with the proposal that the RP is generated by random neural fluctuations in frontal cortical areas, or that it reflects uncertainty in the timing of action. Our findings raise new questions about the neural antecedents of self-initiated actions, and the mechanisms underlying temporal decision-making.

## Supplementary Materials

### Supplementary Behaviour

**Figure S1.**
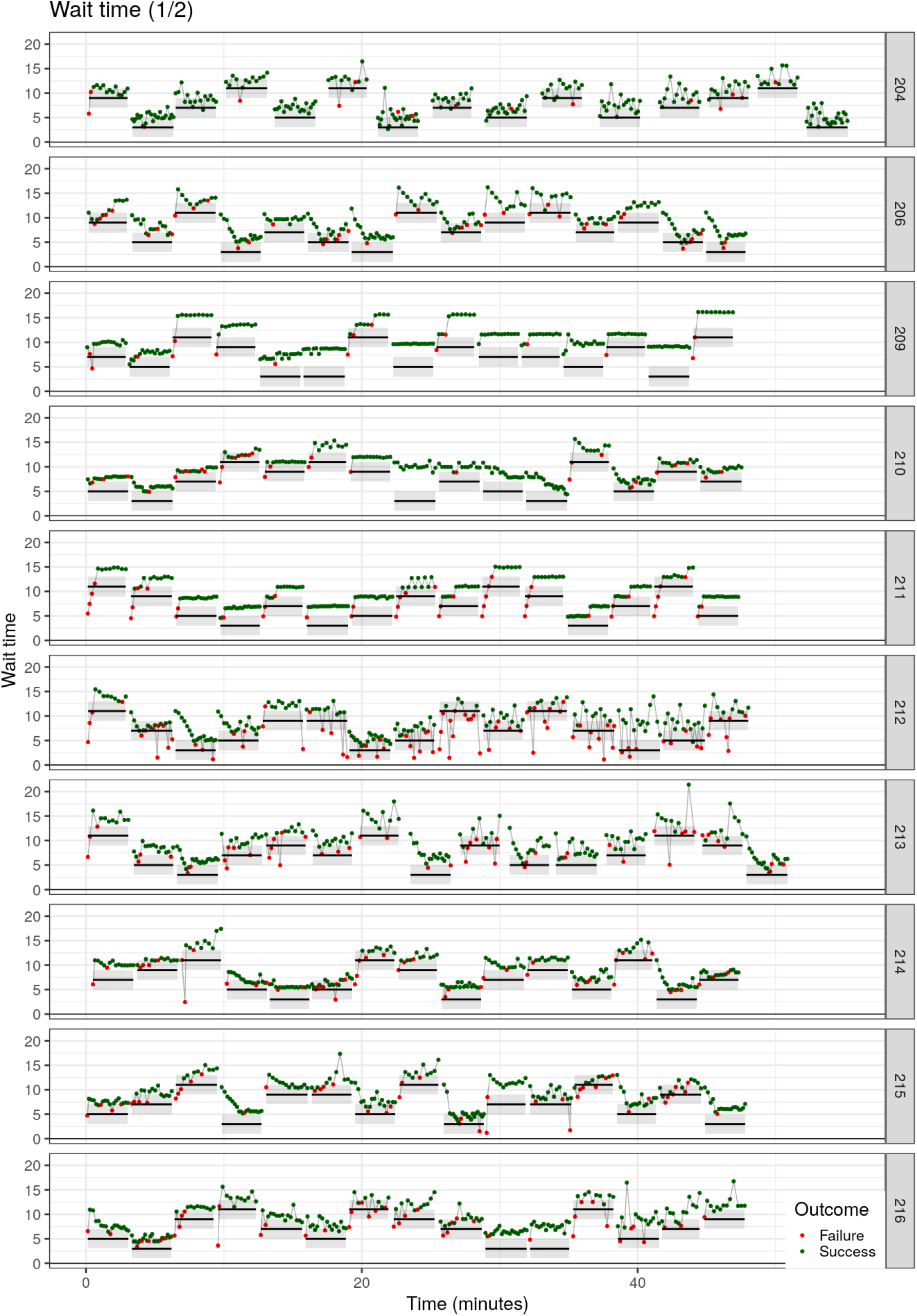
Waiting times across the experiment for individual participants (1 of 2). Horizontal bars show mean ±2SD times at which soufflés were ready in each block. Green (red) dots show trials where participants did (not) wait long enough.

**Figure S2.**
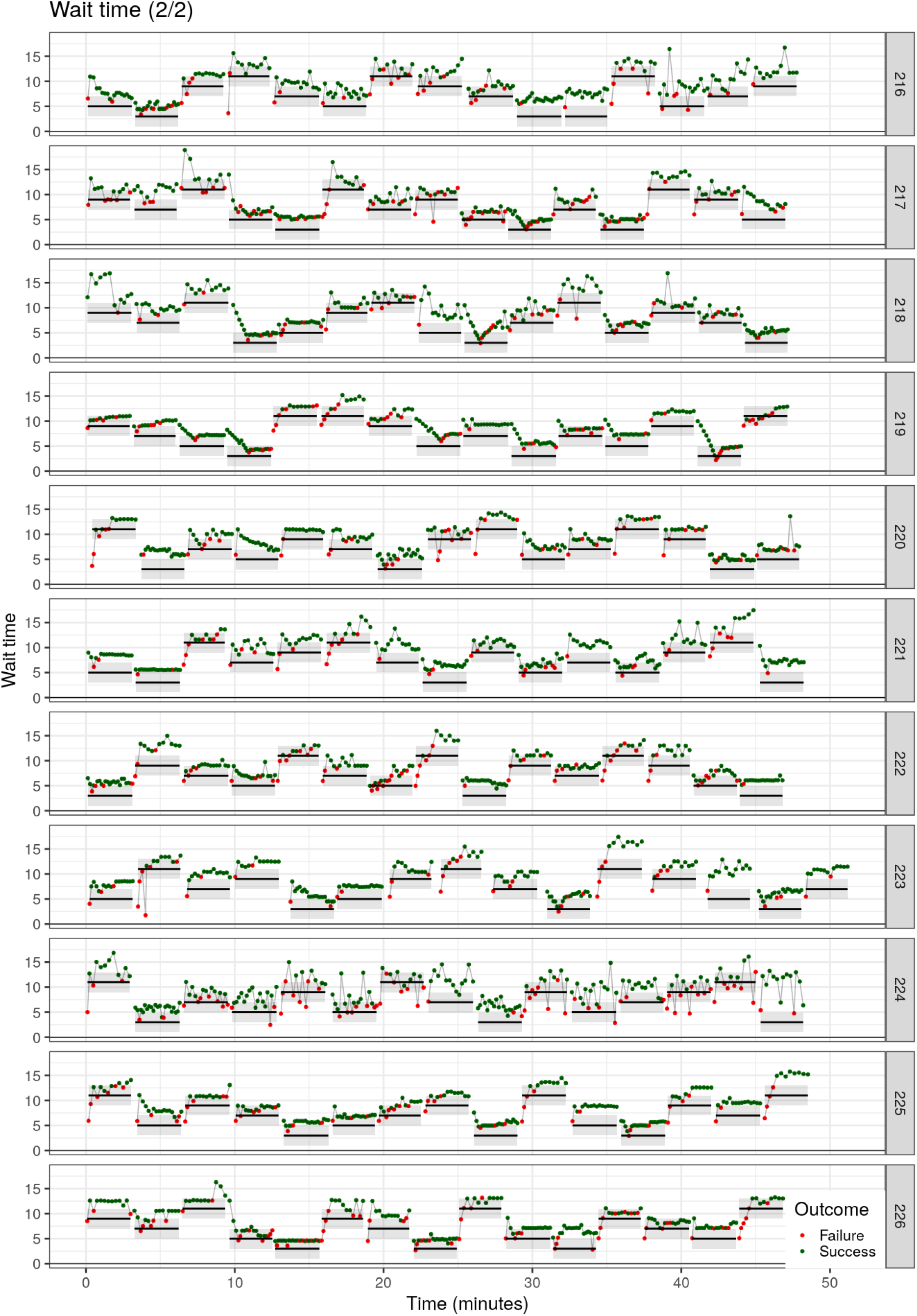
Waiting times across the experiment for individual participants (2 of 2).

**Figure S3.**
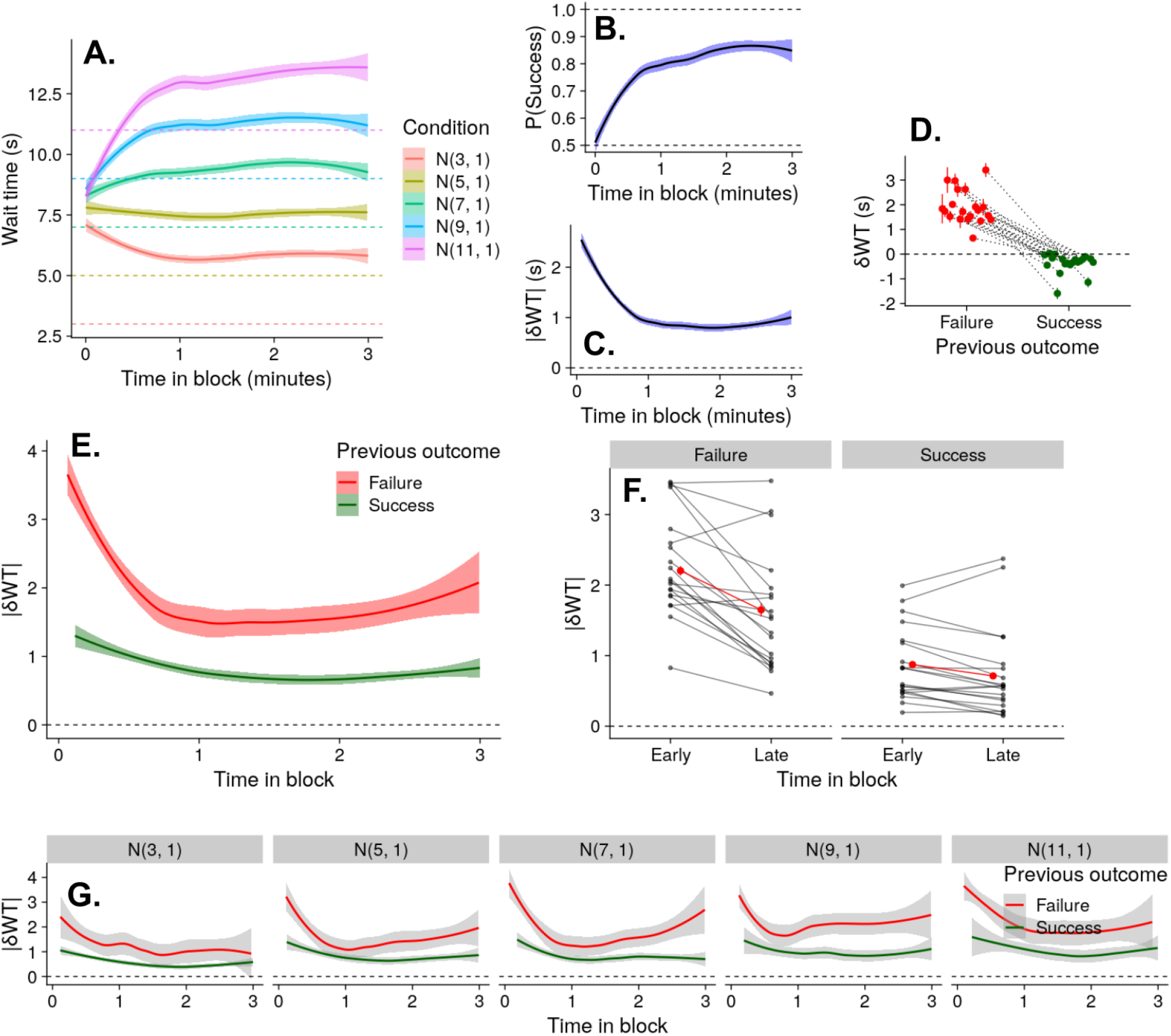
Supplementary behaviour. **A.** Smoothed average wait times over time in each condition. Dashed lines show mean ready times for each condition. In this and subsequent plots, data is pooled across participants, and shaded region shows standard error. “N(3, 1)” denotes the condition where soufflés were ready after 3 seconds on average, with a standard deviation of 1 second. **B.** Probability of successfully waiting until the soufflé is ready, over time. **C.** Absolute change in wait times, over time. **D.** Changes in wait time as a function of the previous outcome. **E.** Absolute changes in wait time as a function of the previous outcome, over time. **F.** Absolute changes in wait time as a function of previous outcome, and time in block (early = first 90 seconds). **G.** Panel E, split by condition.

### Supplementary Analysis: Regression to the Mean

Participants waited considerably longer after trials where they acted too soon (failures) and acted slightly sooner after a previous trial where they waited long enough to retrieve the soufflé (successes), suggesting that they followed some form of reinforcement learning strategy. However, waiting times were also shorter for failures (M = 7.8 s, SD = 0.6 s) than for success (M= 9.1 s, SD = 0.5 s), t(19) = 8.456, p < .001. This raises the possibility that the effect of the previous outcome may be explained by regression to the mean: participants might simply slow down after faster-than-usual actions, and speed up after slower-than-usual actions.

To ensure that regression to the mean alone does not explain the relationship between previous outcomes and wait times, we fit a linear mixed model with δWT (seconds) as the outcome,WT (seconds) as the outcome, and both previous outcome (binary) and previous wait time (seconds) as predictors. There was a significant effect of previous outcome, indicating that participants learned from previous actions, b = −2.14, CI = [−2.25, −2.02], t(4796.7) = 36.4, p < .001, and a significant effect of previous wait time, indicating some regression to the mean, b = −0.16, CI = [−0.18, −0.15], t(4793.4) = 20.0, p < .001.

### Supplementary EEG

**Figure S4.**
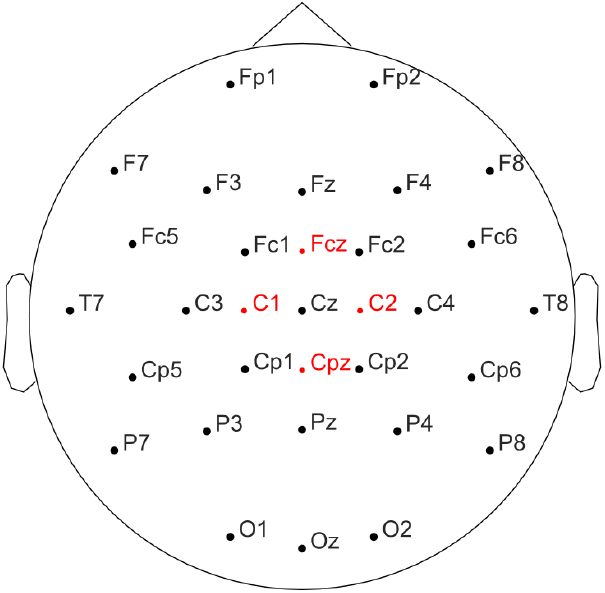
Electrode positions. The four electrodes excluded from the average reference are marked in red.

**Figure S5.**
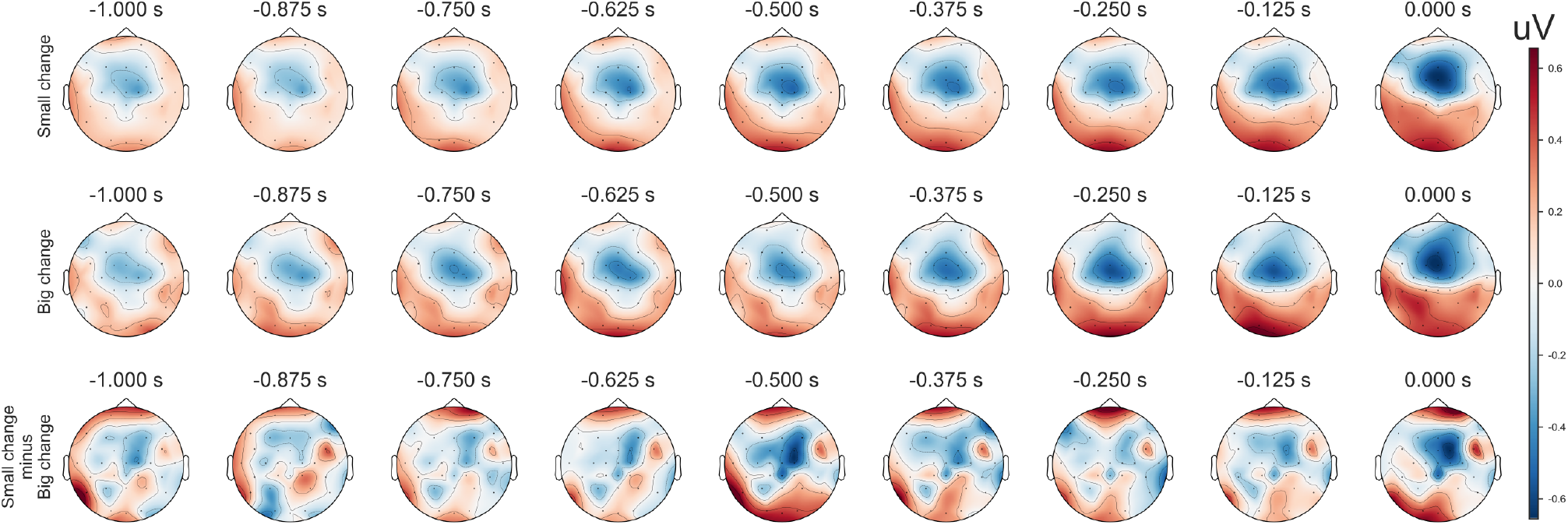
Scalp distributions of the RP in the second prior to action on trials with small absolute changes in wait time (top), large changes (middle), and the difference wave (bottom). The negative RP component centred around electrode FCz is more pronounced for trials with small changes in wait times.

**Figure S6.**
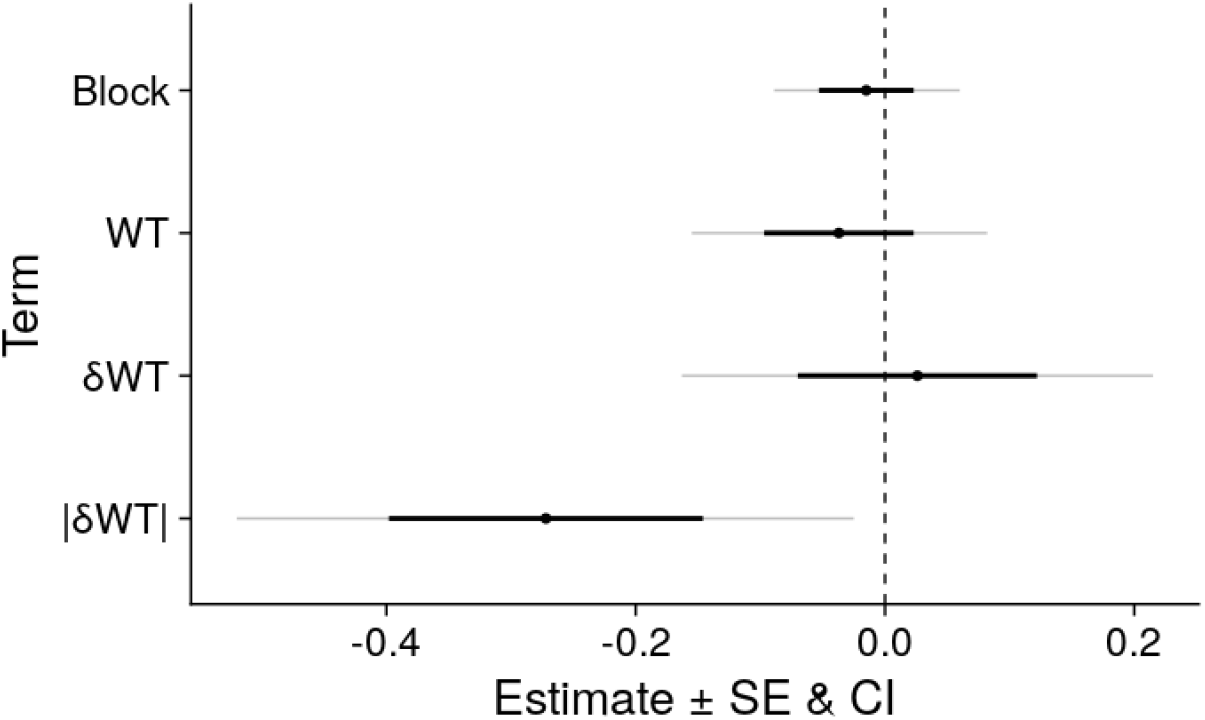
Regression parameter estimates for the linear mixed model reported in the manuscript. Black error bars show SE. Grey bars show 95% confidence intervals. Block = Experimental block (1 to 15, centred as −7 to +7). WT = Wait time in seconds. δWT = Signed change in wait time from previous trial. |δWT| = Absolute change in wait time.

